# Standardization of laboratory practices for the study of the human gut microbiome

**DOI:** 10.1101/2022.11.10.515556

**Authors:** Jolanda Kool, Liza Tymchenko, Sudarshan Shetty, Susana Fuentes

**Author notes:** Corresponding author. National Institute for Public Health and the Environment (RIVM), Antonie van Leeuwenhoeklaan 9, 3721 MA Bilthoven, the Netherlands.

## Abstract

Technical advances in next-generation sequencing (NGS) have made it more accessible to study the human microbiome, resulting in more available data and knowledge. As a result of this expansion of data, the need to obtain comparable and reproducible data has become one of the most important challenges facing microbiome research nowadays. In this study, we aim to contribute to existing knowledge to promote high quality microbiome data and minimize bias introduced by technical variation throughout studies, from sample collection, storage, to sequencing strategies. While immediate freezing upon sampling has been the “golden standard” in the field, this method is often logistically difficult and expensive, becoming a limiting factor when conducting large scale studies or in regions where maintenance of the cold-chain presents difficulties. Therefore, we compared the immediately frozen method to storage at room temperature for 3 – 5 days in two commercially available stabilization solutions (Omnigene gut and Zymo Research) as well as without buffer. Other important aspects were tested, such as DNA extraction, bacterial DNA input or number of PCR cycles. Method choice for cell disruption resulted in the biggest difference in compositional profiles. The changes observed in microbiome profiles in samples stored at RT without stabilization solution was prevented by the use of these. For library preparation and sequencing, we found the highest heterogeneity in the DNA extraction step, followed by the use of different Illumina barcodes, indicating that both of these steps have an impact during library preparation. We did not observe a batch effect between the different sequencing runs. Standardized methods are important to allow comparison of results between different research groups worldwide and reliably expand microbiome data to a broad range of diseases, ethnical backgrounds and geographic locations. A more global perspective will increase our understanding of the human microbiome around the world.

## Introduction

Over the last decades, numerous studies have shown the value of investigating the role of the human microbiome in health and disease. The intestinal microbiome has been associated with several disorders, ranging from gastrointestinal diseases like inflammatory bowel disease or colorectal cancer [1, 2] to systemic disorders such as obesity and diabetes [3–5]. The intestinal microbiome interacts with the host immune system [6, 7], and has been shown to play a role in the response mounted to vaccines, such as vaccine-induced gut mucosal antibody response to the oral polio vaccine (OPV) [8] and the rotavirus vaccine (RVV) [9]. In recent years, technological advances in next-generation sequencing (NGS) have made it more accessible to study the human microbiome. With a reduction in sequencing costs and an increase in capacity for high-throughput sequencing, the number of studies that investigate the human microbiome has expanded dramatically. Although this increase in available data and knowledge is beneficial for the field, the urge to ensure reproducible and comparable results has become one of the most important challenges facing researchers nowadays.

Several studies compare different methods for microbiome research through the entire workflow, from sample collection to sequencing approaches [10, 11]. Recently, the STORMS checklist was published providing a guideline for concise and complete reporting of microbiome studies [12]. For gut microbiome research, the most widely accepted method for sampling of faecal samples is freezing upon collection, in which participants are usually asked to collect their faecal sample at home and store it in a home freezer before transportation to the laboratory for further processing. While optimal for sample storage, this method is often logistically difficult and expensive, becoming a limiting factor when conducting large scale studies, especially in regions where maintenance of the cold-chain during transportation is challenging. Therefore, alternative sampling methods are essential to facilitate large population level microbiome studies from broad geographic locations. To this end, several approaches have been tested aiming to stabilize the microbial composition in faecal samples during transportation at ambient temperature, such as the Omnigene gut system, the Stratec stool collection tube, or the Stool Nucleic Acid Collection and Preservation Tube [13–15].

Furthermore, after sampling, extraction of nucleic acids is shown to be potentially the most critical step affecting the outcome of microbiome studies. Decisions in approaches to break the cell wall, through chemical lysis or mechanical disruption (bead beating), or even composition and size of beads used, can significantly impact the composition of the faecal microbiome[16, 17]. After DNA is extracted, several other factors like PCR conditions, choice of primers and downstream bioinformatics approaches can, although to a lesser extent, impact the end results of faecal microbiome analyses [18,19].

In our study, we aimed to contribute to the existing knowledge and work towards a controlled and reproducible wet-lab workflow for the study of the faecal microbiome. To that end, we examined the effect of different sample collection and short-term storage conditions, using the Omnigene gut and Zymo research stabilization systems. In addition, we studied the effect of cell disruption using chemical or mechanical methods, and tested different DNA extraction kits. Finally, we investigated the impact of bacterial input and number of PCR cycles during library amplification, and analysed the inter- and intra-run variations in large scale studies, by testing the effect of DNA extraction rounds and the use of different barcodes on the overall microbial composition.

## Material and methods

### Sample selection and study design

Samples were selected from the PIENTER3 and Z-test study (Table 1). PIENTER is a cross-sectional study, designed to periodically monitor the seroprevalence of National Immunization Program (NIP)-targeted diseases in the Netherlands. During the third survey in 2016-2017 (PIENTER3), faecal samples were collected from participants throughout the Netherlands and the Caribbean islands Bonaire, St. Eustatius and Saba [20]. The Z-test study was designed for protocol optimization. None of the participants of both studies had undergone antibiotic treatment in the three months prior to collection and informed consent was obtained from all participants.

**Table 1.**
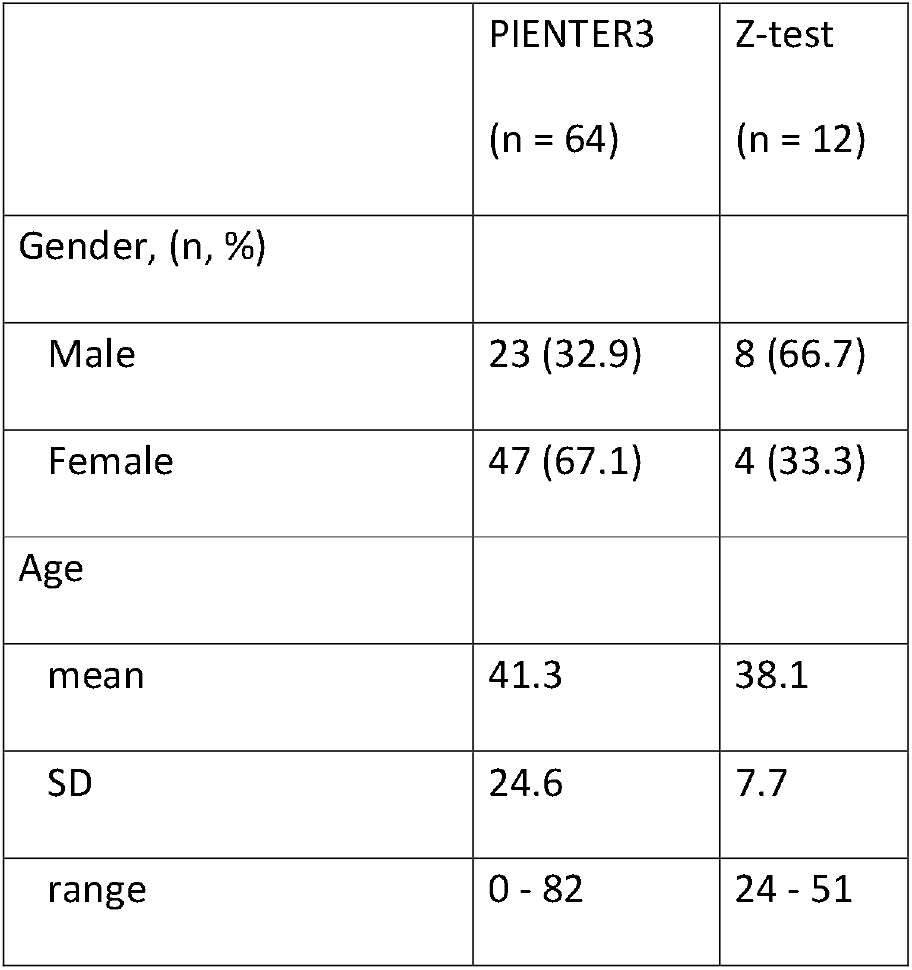
Demographics of the study participants.

|  | PIENTER3<br>(n = 64) | Z-test<br>(n = 12) |
| --- | --- | --- |
| Gender, (n, %) |  |  |
| Male | 23 (32.9) | 8 (66.7) |
| Female | 47 (67.1) | 4 (33.3) |
| Age |  |  |
| mean | 41.3 | 38.1 |
| SD | 24.6 | 7.7 |
| range | 0 - 82 | 24 - 51 |

To investigate the effect several aspects known to impact a microbiome study (e.g. sample collection, nucleic acid extraction, library preparation and sequencing through different approaches (Fig. 1)), we compared different approaches and results were analysed using 16S rRNA gene amplicon sequencing data.

**Figure 1.**
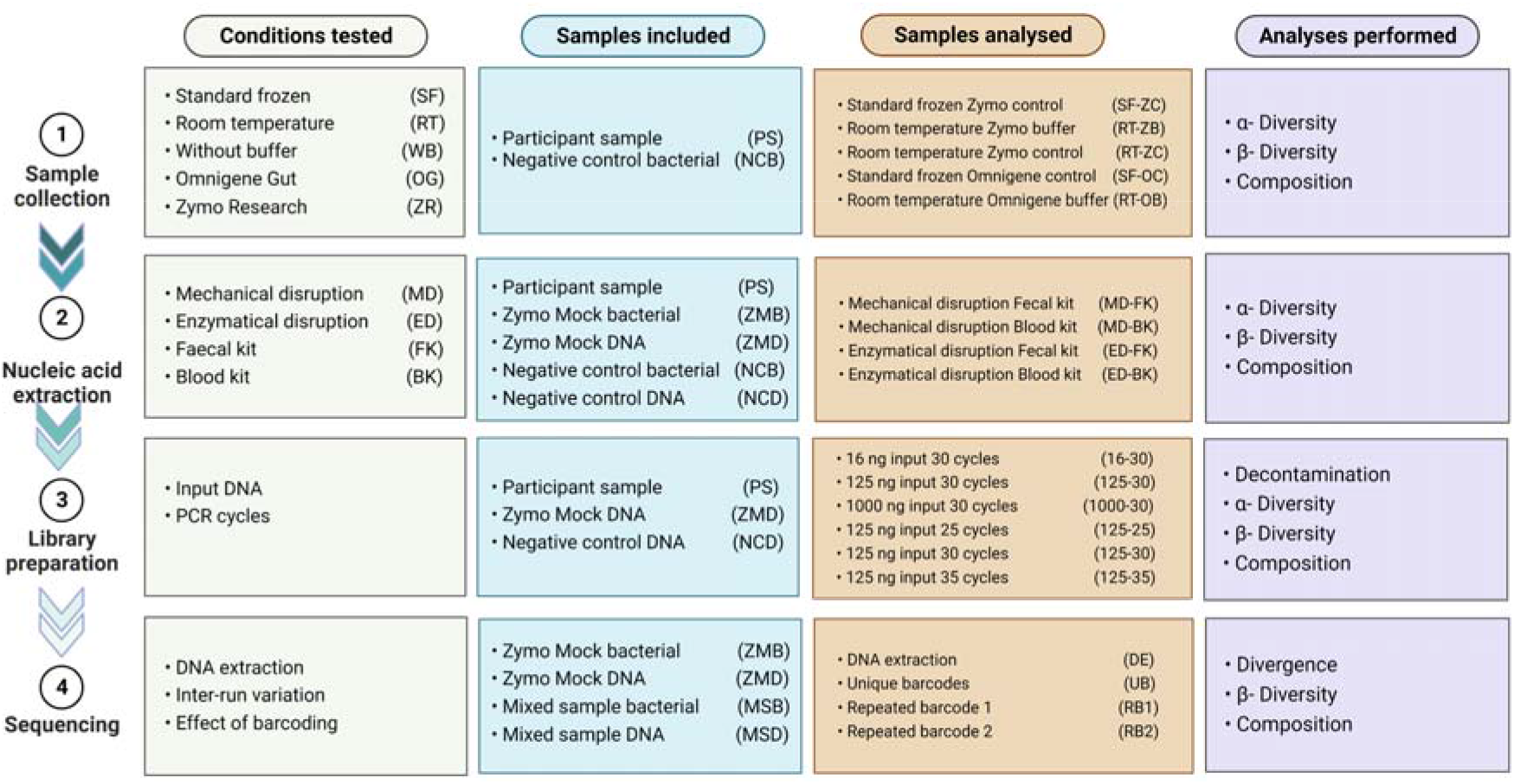
Overview of the conditions tested, samples included and processed in every experiment and analyses performed.

### Sample collection

For sample collection under standard conditions, participants were asked to freeze their faecal samples in their home freezers (approximately −20°C) directly upon collection. Subsequently, samples were transported to the laboratory on dry ice and stored at −80°C until further processing. Within the PIENTER3 study, a subset of participants (n=64) collected samples in duplicate. Next to the standard condition, samples were collected in the Omnigene gut tube (DNA Genotek, Ottawa, Canada) containing 2 mL stabilization buffer and stored at room temperature (RT) for 3-5 days before freezing (−80°C). For the Z-test study, all participants (n= 12) collected samples in triplicate. The first sample was collected under standard conditions, the second sample was stored at RT for 3 – 5 days and the third sample was collected in a Zymo research tube (Zymo Research Inc., Irvine, CA, USA) containing 9ml DNA/RNA shield and stored at RT for 3 – 5 days before storage at −80 °C. A positive control (mixed sample-bacterial or “MSB”) was generated by thoroughly mixing equal amounts of faecal material from 5 randomly selected healthy participants. This mix was evenly aliquoted and stored at −80°C until further use in different random DNA extraction runs (n= 23). DNA extracted from these aliquots (mixed sample-DNA or “MSD”) was used as control in different library preparations (n = 12). In addition, microbial community standards (ZymoBIOMICS, Zymo Research, Irvine, CA, USA) were used as positive controls. We used the ZymoBIOMICS Microbial Community Standard as control for DNA extractions (ZMB) and the ZymoBIOMICS Microbial Community DNA Standard as control during library preparation (ZMD). Blank negative controls were included in every DNA extraction, where the appropriate buffer was used depending on the collection method, without adding any faecal material. DNase-free water was used as blank during library preparation.

### DNA extraction

DNA was extracted by mechanical disruption of the cells (n= 358), and a subset (n= 24) were in parallel treated via enzymatic lysis. To lyse the cells enzymatically, tubes containing 0.25g of faecal material, 1 mL of Lysis buffer (Promega, Madison, USA) and 40 μl of Proteinase K (Promega, Madison, USA) were vortexed and heated at 95°C for 5 min. After heating, the samples were cooled down and incubated at 56°C for 5 minutes. Samples were centrifuged (17,000xg for 5 minutes) and the supernatant lysates were collected for further processing. For mechanical disruption, we used a repeated bead-beating and column purification process, using pre-assembled tubes containing 0.5g zirconia/silica beads (0.1 mm) and 5 glass beads (2.7 mm) (Biospec products, Bartlesville, OK, USA). In tubes without stabilization buffer, 0.25g of faecal material and 700μl S.T.A.R. buffer were added to the beads. For faecal material collected in the Zymo research tubes, 1 mL of the stabilization buffer containing the faecal sample was added to the beads. From the material collected in the Omnigene Gut tubes, 250 μl of the stabilization buffer and 450μl S.T.A.R. buffer were added to the beads. All samples were lysed by repeated bead-beating (5,5 ms for 1 minute, repeated 3 times and cooled on ice in between) in a Fastprep-24 (MP Biomedicals, Irvine, USA), followed by heating of the samples at 95°C for 15 min. Samples were centrifuged and the lysates collected. All lysates were further purified in the Maxwell RSC instrument (Promega, Madison, USA), using the Maxwell RSC blood DNA kit (n=376), and the Maxwell RSC Fecal kit on a subset of the samples (n= 28). DNA was eluted in 60 μl elution buffer and further purified by using the OneStep PCR Inhibitor Removal Kit (ZymoBIOMICS, Zymo Research, Irvine, CA, USA).

### Quantification of bacterial DNA by quantitative PCR

DNA concentration was measured using a Quantus fluorometer (Promega, Madison, USA) and samples were stored at −20°C until further processing. The bacterial load present in the purified samples was measured by quantitative PCR (StepOnePlus Real-Time PCR System, Thermo Fisher Scientific, the Netherlands), using a universal primer set targeting the 16S rRNA gene (forward Eub34lF: CCTACGGGAGGCAGCAG, reverse Eub534R: ATTACCGCGGCTGCTGGC) [21]. The quantitative PCR was carried out by using SYBR Green, in a 25 μl reaction consisting of 12,5 μl Maxima SYBR Green/ROX qPCR Master Mix (ThermoFisher scientific, Waltham, MA, USA), 0,5 μM forward primer Eub341F, 0,5 μM reverse primer Eub534R, 2 μl DNA (500 times diluted in HPLC grade water) and 8 μl HPLC grade water. DNA was denaturated (95°C; 10 min), followed by 40 cycles of denaturation (95°C; 15 sec), annealing (60°C; 15 sec), extension (72°C; 15 sec) and a holding stage (95°C; 1 min and 60°C; 1 min).

### Library preparation and sequencing of the V4 region of the 16S rRNA gene

Results obtained from the qPCR were used to equalize the amount of bacteria present in all samples and provide an input of 100 pg DNA for amplification of the hypervariable V4 region of the 16S rRNA gene, using the 515F (5’-GTG CCA GCM GCC GCG GTA A-3’) and 806R (5’-GGA CTA CHV GGG TWT CTA AT-3’) primers, including the Illumina flow cell adapter and a unique 8-nt index key [22–24]. The amplification mix consisted of 0,5 μl (1 U) Phusion Hot Start II High-Fidelity DNA Polymerase, 5 μl 5x Phusion HF Buffer (Thermo Fisher Scientific), 7 μl HPLC grade water, 2,5 μl of 2mM dNTP mix (ThermoFisher scientific, Waltham, MA, USA), 0,5 μM of forward primer 515F, 0,5 μM of reverse primer 806R and 5 μl template DNA. After denaturation (98°C; 30 sec), 30 cycles were performed consisting of denaturation (98°C; 10 sec), annealing (55°C; 30 sec), extension (72°C; 30 sec) and a final hold (72 °C; 30 sec). For optimization of the V4 amplicon PCR, the effect on the microbiota profile using a different amount of input DNA (16, 125 and 1000 pg) and the number of cycles (25, 30 and 35 cycles) were investigated using 42 samples. Amplified product was checked on size and quantified to pool equimolar, using the QIAxcel DNA High Resolution Kit on the Qiaxcel Advanced System (Qiagen, Hilden, Germany). The pool was purified twice by 0.9x AMPure XP magnetic beads (Beckman Coulter, the Netherlands). Final quantification of the pool was done using the KAPA library quantification kit (Roche, USA). Paired-end sequencing was conducted using a V3 Miseq reagent kit (600 cycles) on an Illumina Miseq instrument (Illumina Inc., San Diego, CA, USA).

### Processing of 16S rRNA gene sequence data

Sequence data was demultiplexed based on sample specific barcode combinations and the primers were removed, prior to processing of the raw reads using the DADA2 pipeline. Default parameters were used unless otherwise stated [25]. Reads were trimmed at 220nt and 100nt for forward and reverse reads respectively, and filtered by truncating reads of a quality score less than or equal than 5. For inferring sequence variants, a minimum of 1E^9^ bases were used for error rate learning. The resulting amplicon sequence variants (ASV’s) were cleared from chimera’s and taxonomy was assigned with the RDP classifier and SILVA database (version 138.1) [26, 27].

### Statistical analysis

Analyses were performed in R version 4.1.0 [28]. Alpha-diversity indices and beta-diversity ordinations (of Bray-Curtis distance) were calculated with the “phyloseq” package [29]. Smaller datasets (n= 76) were visualized with nonmetric multidimensional scaling (NMDS), while PCoA was used for larger datasets. To determine the differences in the Alpha-diversity measures, the Wilcoxon test from the rstatix package was used, and corrected for multiple testing using the Bonferroni method where appropriate. Community level differences in beta-diversity were tested using PERMANOVA adonis function from the vegan (v2.5-6) R package.

Linear discriminant analysis effect size (LEfSe [30]) were performed with an alpha-value for the factorial Kruskal–Wallis test among classes of >0.05 and a LDA threshold of >4.0. Results were visualized in a cladogram. Identification of contaminant ASV’s was done with the decontam (v1.8.0) R package using the prevalence method (threshold = 0.1) [31]. Correlations of the Mock samples compared to the theoretical composition were calculated with the Spearman’s correlation using the “checkZymoBiomics” function from the chkMocks (v0.1.03) package [32].

## Results

### Effect of sample collection on the overall diversity and composition of the faecal microbiome

To investigate the impact of method choice for sample collection on the overall microbiome diversity and composition, faecal material of participant samples (PS) and negative bacterial controls (NCB) were used. Preservation of faecal material at room temperature (RT) was tested in three conditions, tubes with Zymo Research buffer (ZB), Omnigene Gut buffer (OB) and a control sample without buffer (ZC and OC), and compared to sample storage under standardly frozen conditions (SF) (Fig. 2).

**Figure 2.**
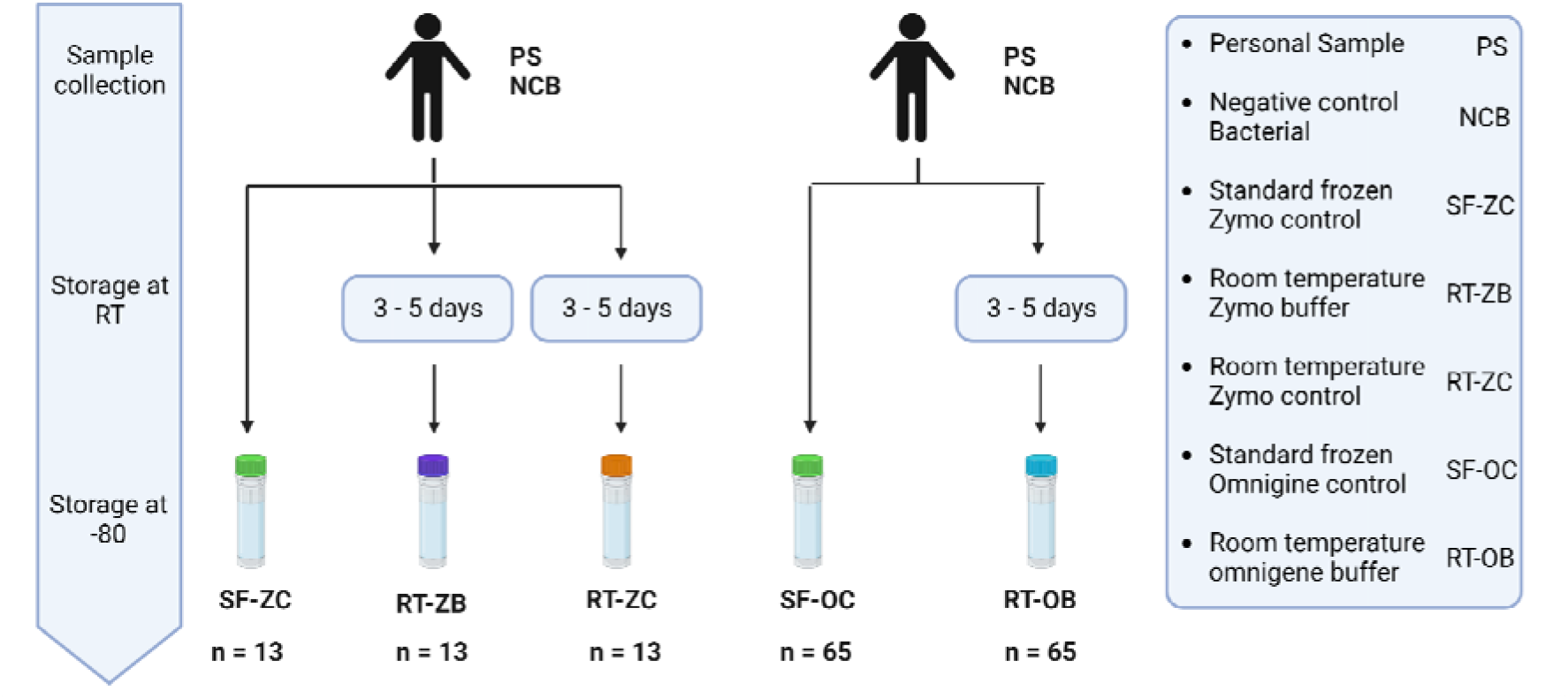
Overview of the different methods for sample collection and storage. The effect of storage at RT for 3-5 days was tested with and without stabilization buffer of the Zymo research (ZB) and Omnigene Gut (OB) collection tubes and compared to the standard storage condition (SF). Faecal material of 12 participant samples (PS) and one negative bacterial control (NCB) was tested in triplicate: one control sample stored under standard storage conditions (SF-ZC), and two additional copies stored at room temperature for 3-5 days prior to freezing at −80 °, with and without the Zymo Research stabilization buffer (named RT-ZB and RT-ZC respectively). Additionally, faecal samples of 64 participants (PS) and one negative bacterial control (NCB) were collected in duplicate. Besides the control sample collected under standard conditions (SF-OC), a copy was collected in the Omnigene gut tube with a stabilization buffer and stored at RT for 3-5 days before freezing at −80 ° (RT-OB).

No differences between the different collection and storage conditions were observed in alpha diversity measures, as calculated using the Shannon index, Observed taxa and Simpson’s indices (Supplementary Fig. 1a). Next, we looked at the overall differences in microbial composition by using the Bray-Curtis distance in a PCoA ordination (Fig. 3a). The largest differences were driven by the two different study groups (R^2^ = 0.026; p. adjusted = 0.001). Therefore, to control for this confounding effect, comparisons were performed within the duplicates of each study. Although there was no significant impact on the microbial composition by sample collection in the Zymo Research tubes, a significant effect was observed when the Omnigene gut tubes were used (R^2^ = 0.020, p. adjusted = 0.002). We further investigated the microbial composition at the phylum level (Fig. 3b, Suppl. Fig. 1b). When comparing the stabilization buffers to the standard collection, we observed an overrepresentation of *Bacteroidota* (RT-ZB p= 0,000488, RT-OB p= 2,32E-09) and *Proteobacteria* (RT-ZB p= 0,00342, RT-OB p= 2,91E-06) and underrepresentation of *Actinobacteria* (RT-ZB p= 0,000488, RT-OB p= 7,87E-05), with differences more marked when using the Omnigene gut system. Furthermore, a lower proportion of *Firmicutes* was observed in samples collected using the Omnigene gut collection tubes (p= 0.0025) when compared to the standard frozen condition.

**Figure 3.**
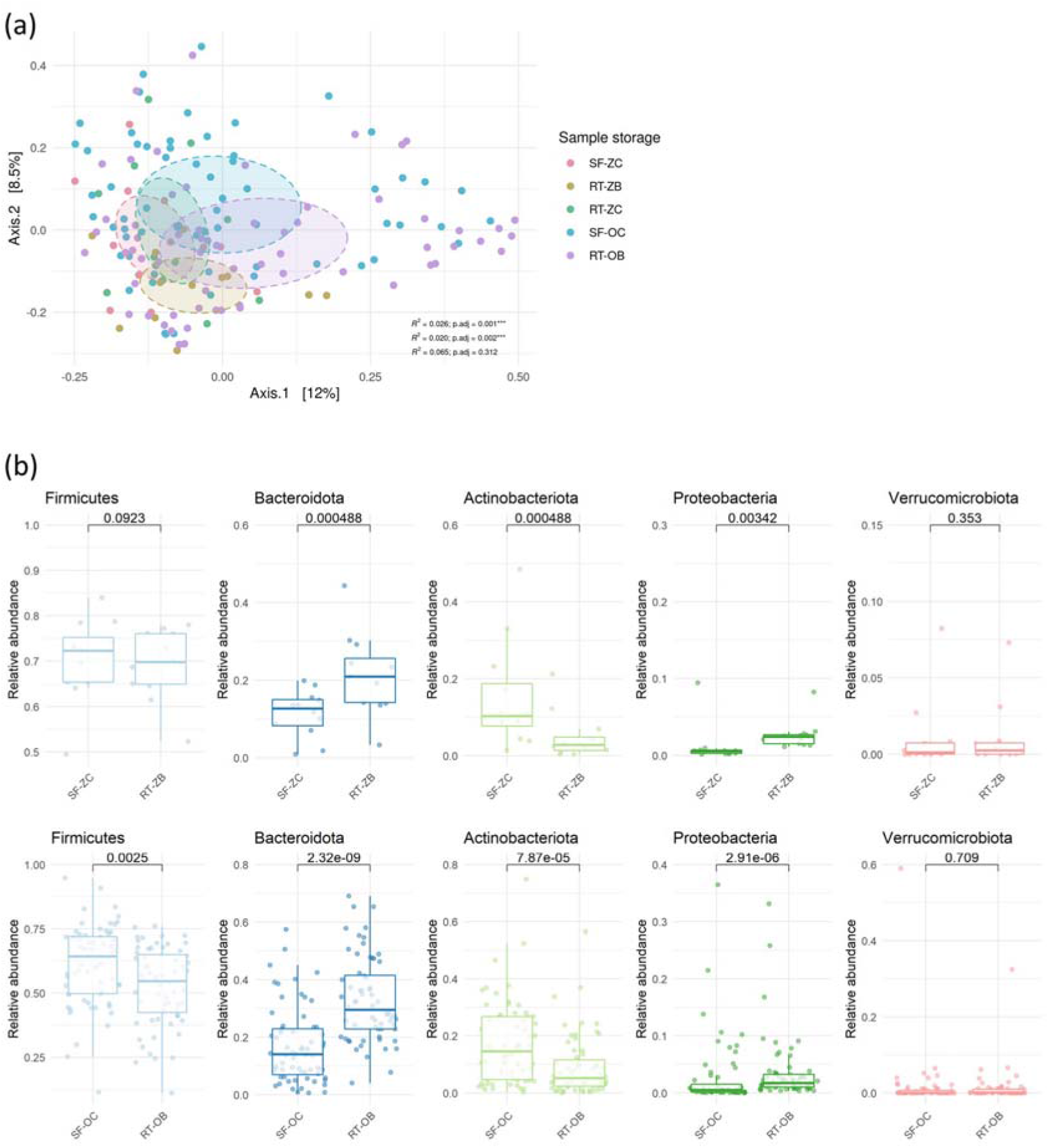
(a) Beta diversity on sample collection and storage. Bray-Curtis distance in a PCoA ordination showed a limited effect of sample collection, only significant for samples stored in the Omnigene gut tubes. perMANOVA testing was performed to test the difference between the two studies (R^2^= 0.026; p. adjusted= 0.001), and the type of stabilization buffer used Omnigene gut tubes (R^2^= 0.020; p.adjusted= 0.002) and Zymo research tubes (R^2^= 0.065; p.adjusted= 0.312). (b) Boxplots show the relative abundance of the 4 top most abundant phyla. The differences between the two studies was tested with the Wilcoxon test.

Specific taxa associated with each storage method were identified using the linear discriminant analysis effect size (LEfSe)(Supplementary Fig. 2, Fig. 4). The taxa that are more abundant in the frozen conditions are represented in red, (SF-ZC, SF-OC) and in green those taxa found more abundant when compared to samples stored with or without stabilization buffers at RT (Fig. 4a-c for comparisons with RT-ZB, RT-ZC and RT-OB respectively). We observed a higher amount of the *Escheríchia-Shigella* group (LDA= 4,63) in samples stored at RT without stabilization buffer when compared to the frozen samples, which was not observed in storage with stabilization buffer. As previously observed, with the RT-OB several taxa within the *Firmicutes* and *Actinobacteriota* were less abundant, including *Blautia* (LDA= 4,33) and *Bifidobacterium* (LDA= 4,51), while taxa from the *Bacteroidota*, such as *Bacteroides vulgatus* (LDA= 4,07), was present in higher abundance (Fig. 4a). For the RT-ZB, similar differences were observed, although the effect was less pronounced than that observed with the RT-OB (Fig. 4c).

**Figure 4.**
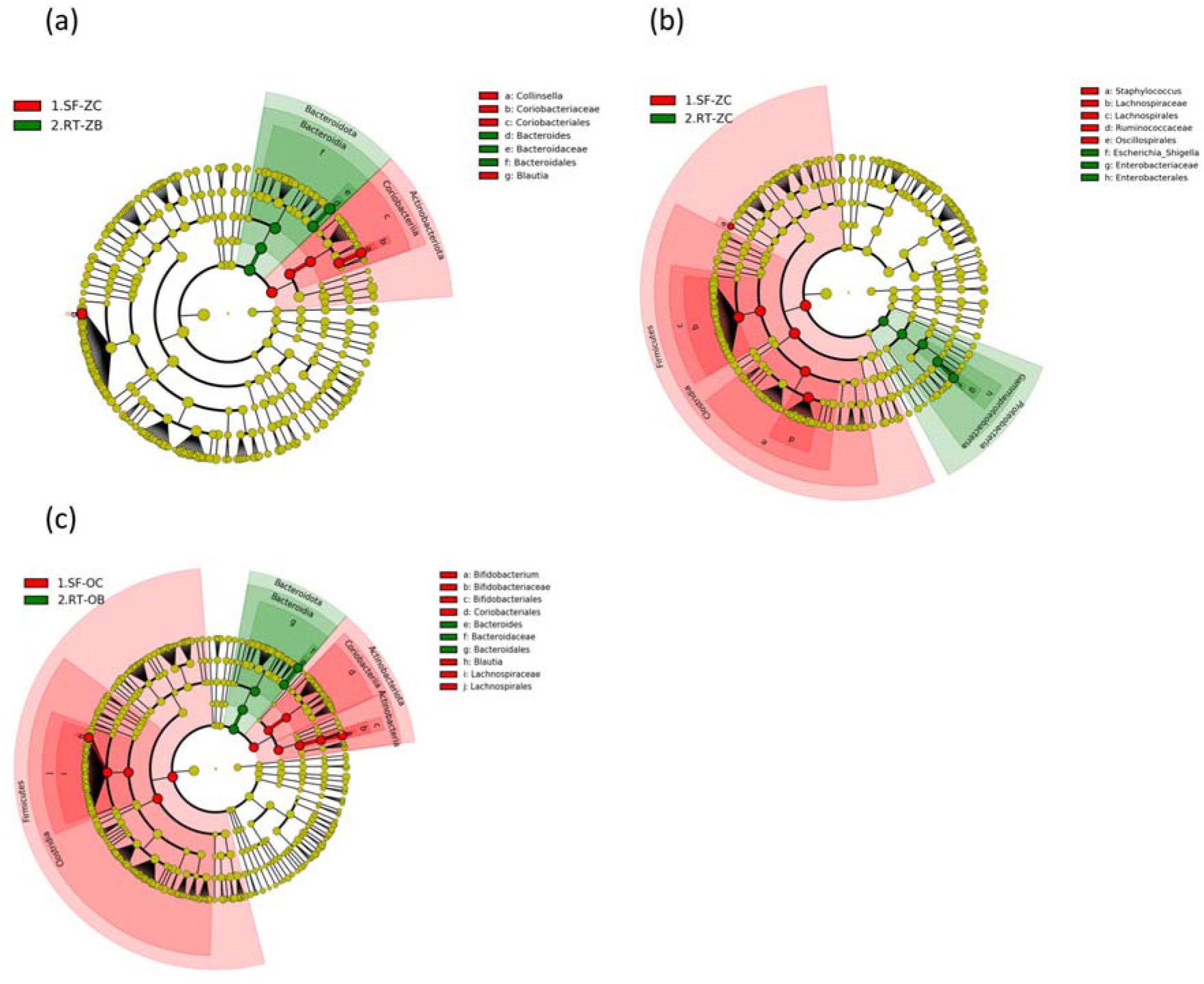
LEfSe analysis showing the significantly distinguishing taxa between the different storage methods based on an LDA score > 4.0. Results are shown in cladograms, showing the effect of storage at RT, with or without stabilization buffer (RT-ZB, RT-ZC, RT-OB) in green, compared to the immediately frozen control samples (SF-ZC, SF-OC) in red. The Cladograms show the taxonomic levels represented by rings, with the phylum level in the outermost ring and the genus level in the innermost ring. Each green or red circle represents a significantly different taxa associated with one of the compared groups.

### Impact of nucleic acid extraction method on the microbial composition

To investigate the effect of the different extraction methods on the faecal microbiome composition and diversity, personal material of 5 different donors, a mock community sample and a negative control was used (Fig. 5). The personal samples of 5 different donors were stored in duplicate, one control sample stored under standard storage conditions (SF-ZC), and an additional copy was stored in the presence of Zymo Research stabilization buffer for 3 – 5 days prior to freezing at −80°C (RT-ZB). The performance of each extraction method was determined by measuring the DNA yield using Quantus fluorometer, and total bacterial yield using a universal 16S rRNA gene qPCR.

**Figure 5.**
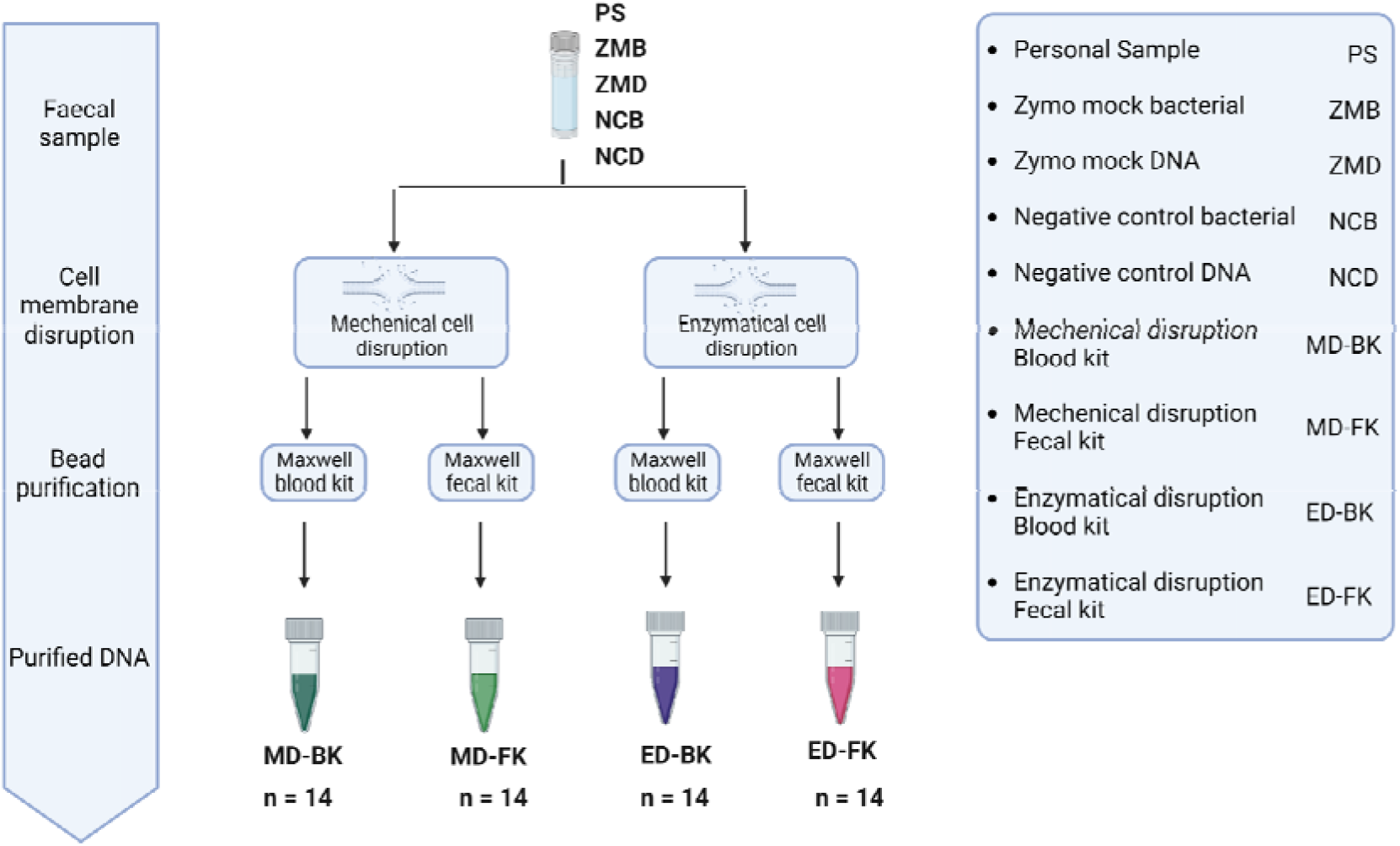
Overview of the different methods for DNA extraction. Personal samples of 5 donors were stored under two conditions, one copy was directly frozen (SF-ZC), the other sample was stored at RT for 3 – 5 days in Zymo research collection tubes (RT-ZB). These feacal samples, together with a positive and negative control were used to test the effect of mechanical or enzymatical cell disruption. Furthermore, we looked into the difference of DNA purification using the Maxwell^®^ RSC Whole Blood DNA Kit and the Maxwell^®^ RSC Faecal Microbiome DNA Kit.

Samples stored without stabilization buffer were extracted more efficiently with mechanical disruption (MD) when compared to enzymatical lysis (ED) (Fig. 6). Interestingly, this effect was not observed in samples stored with stabilization buffer. For these, the best performing method of those tested regarding total bacterial yield, was extraction with the Maxwell RSC Fecal Microbiome DNA kit (FK) when compared to the Maxwell RSC Blood DNA kit (BK), regardless of how the samples were disrupted.

**Figure 6.**
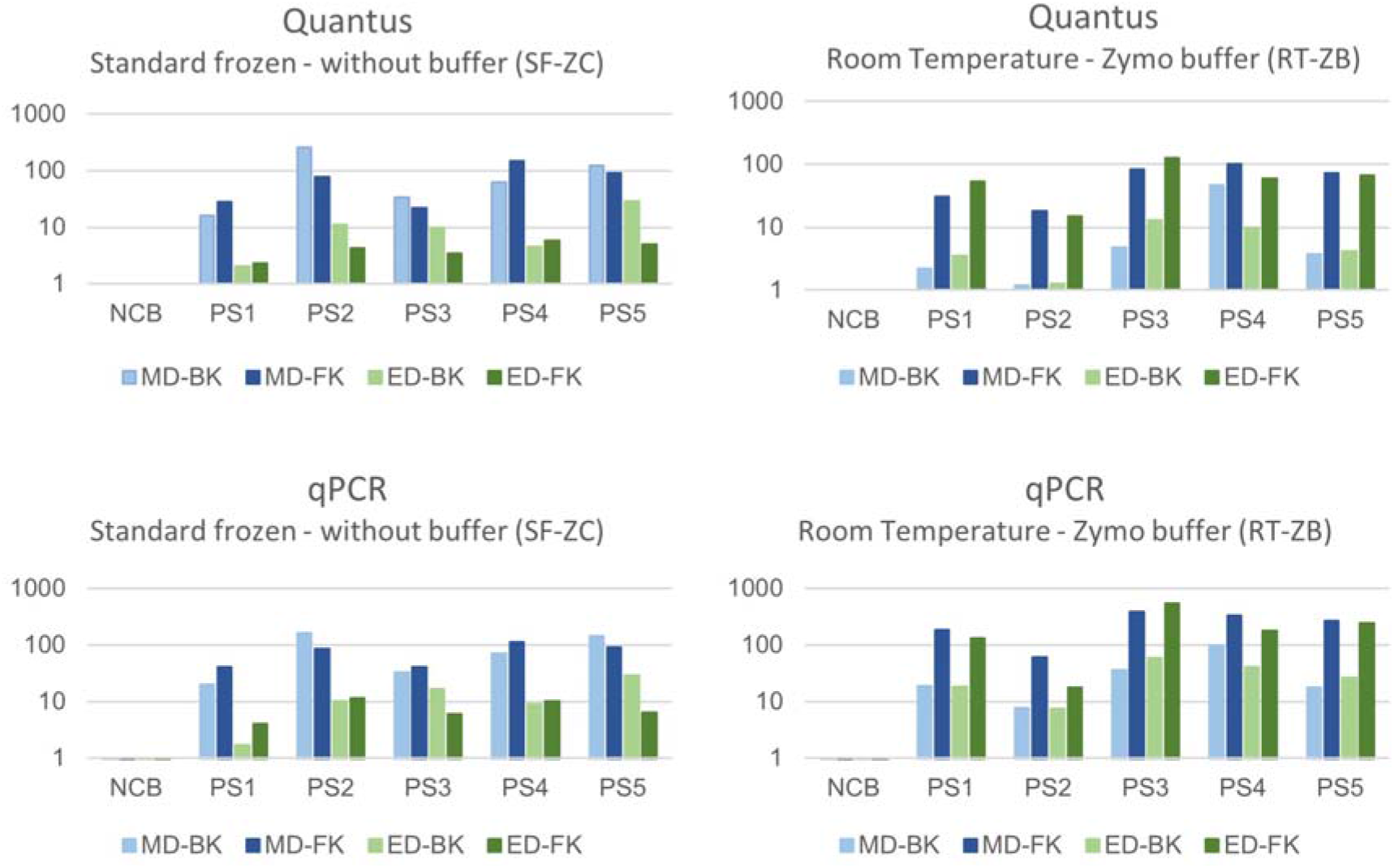
Bacterial yield of samples using mechanical (MD) or enzymatical (ED) disruptionand extracted with the Maxwell RSC Blood DNA kit (BK) or the Maxwell RSC Fecal Microbiome DNA kit (FK). The DNA concentration was measured using the Quantus fluorometer and the bacterial DNA using a universal 16S rRNA gene qPCR and represented in ng/μl.

There were no significant differences in alpha diversity measures between the different extraction methods (Supplementary Fig. 3). On the overall community structure, significant differences were observed between the different methods for cell disruption, however the choice of DNA extraction kit showed no effect on the microbial community (Fig. 7).

**Figure 7.**
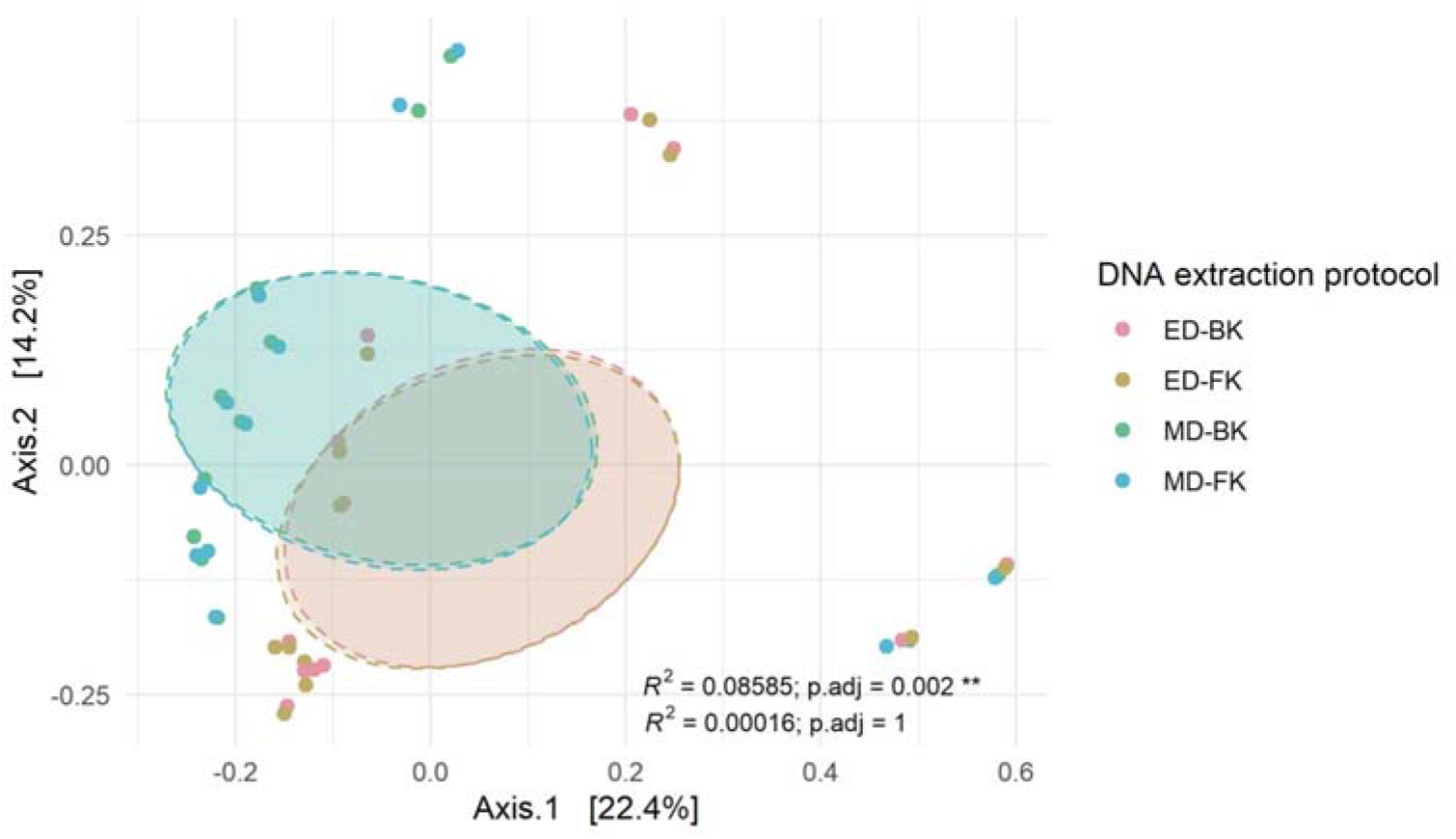
Bray-Curtis distance in a PCoA ordination showing the difference in overall microbial community structure of the different groups (Bead-beating using the Blood and Fecal kits, i.e. MD-BK and MD-FK, and similarly for lysis buffer, i.e. ED-BK and ED-FK). perMANOVA testing for the method for cell disruption (R^2^= 0,08585; p.adjusted= 0,002) and extraction kit (R^2^= 0,00016; p.adjusted= 1)

At the phylum level, no differences were observed between the different DNA extraction kits tested (Fig. 8a). In contrast, the method for cell disruption influenced the microbial composition significantly. Samples that were enzymatically lysed showed an underrepresentation of *Firmicutes* and *Actinobacteriota* and an overrepresentation of *Bacteroidota*, when compared to the mechanically lysed cells.

**Figure 8.**
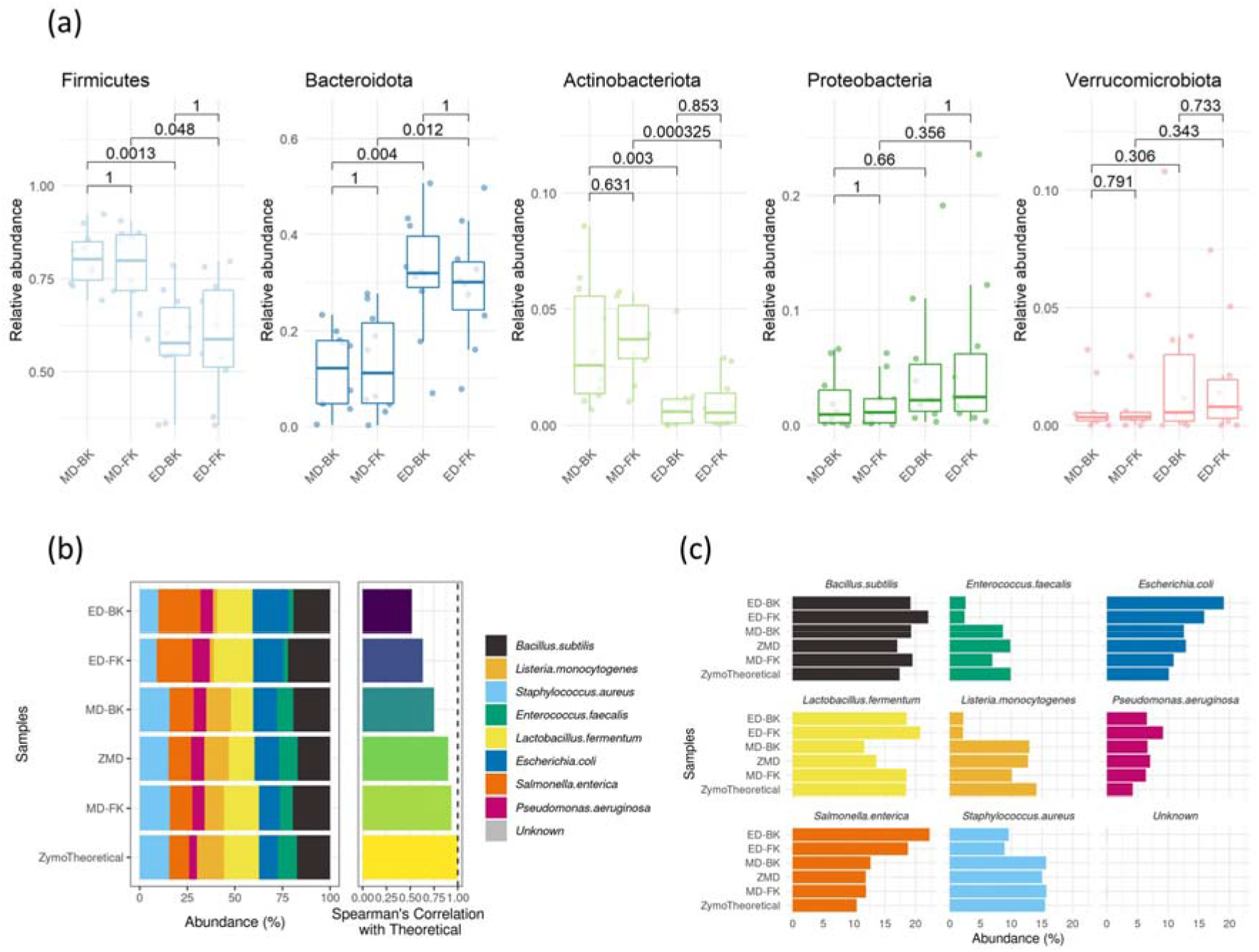
(a) Boxplots show the relative abundance of the top most abundant phyla. The Wilcoxon test was used to calculate the differences between the different DNA extraction methods. (b) Spearman’s correlation of the Mock samples extracted by the different methods, compared to the theoretical composition of the Mock community sample. (b) Barplots of the 8 bacterial strains included in the Zymo Mock sample.

When investigating the mock community sample, by using Spearman’s correlation to the theoretical composition provided by the manufacturer (ds1706_zymobiomics_microbial_community_standards_data_sheet.pdf (zymoresearch.com) we observed the highest correlation in those samples disrupted with bead-beating and lysates purified using the Maxwell RSC Fecal Microbiome DNA kit (rho= 0,933 and 0,633 for MD-FK and ED-FK respectively). The lowest correlation was found in the enzymatically lysed samples, mostly driven by the underrepresentation of the theoretical *Enterococcus faecalis* and *Listeria monocytogenes* (Fig. 8b).

### Limited effect of library preparation on the overall community structure

To assess the effect of differences in protocols during amplification of the V4 region of the 16S rRNA gene, faecal material of 3 different donors and several controls (positive and negative) were used (Fig. 9). We analysed the impact of bacterial DNA input (using 16, 125 and 1000 pg) and PCR cycles (25, 30 and 35 cycles).

**Figure 9.**
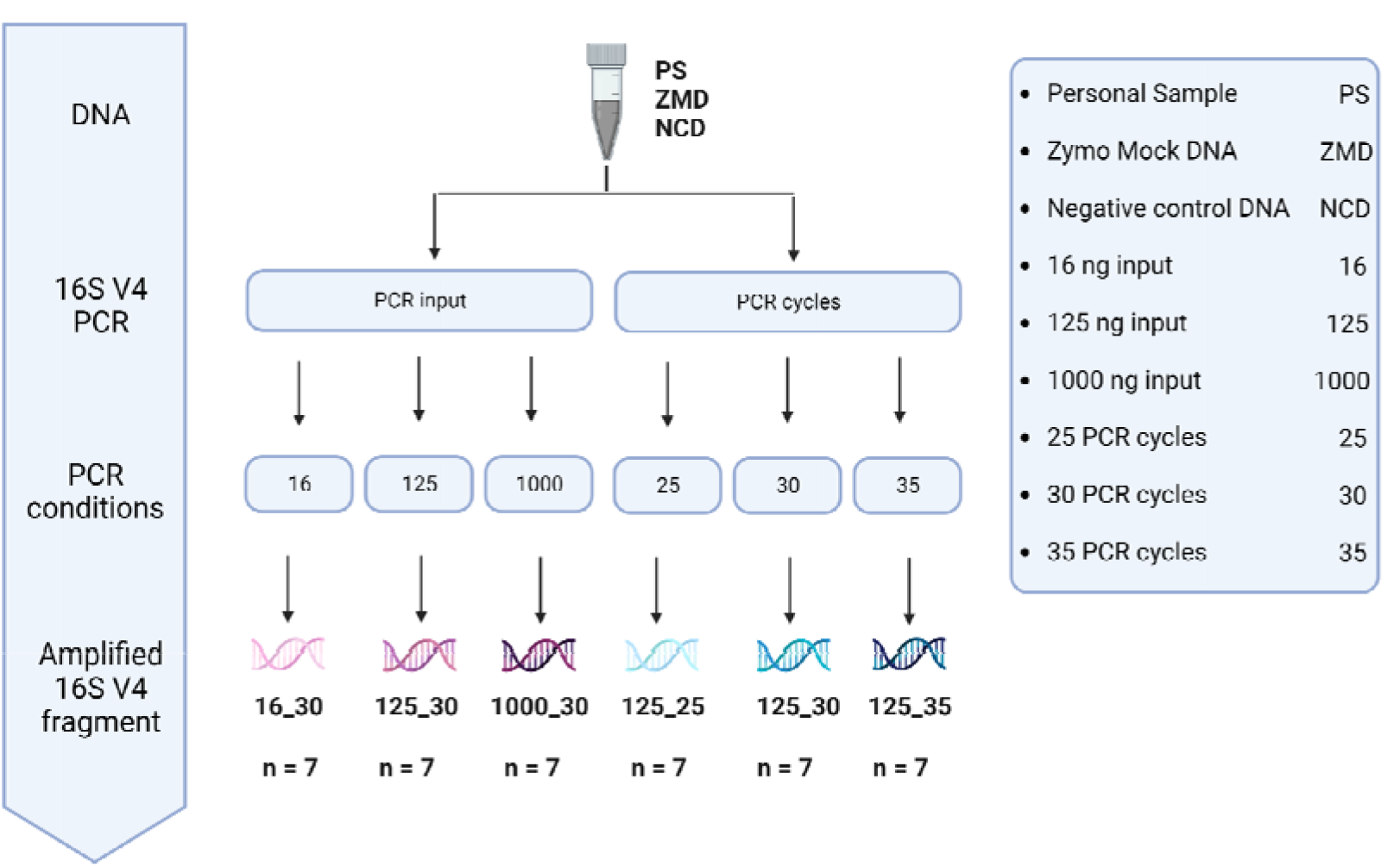
Overview of the different methods tested during library preparation. The effect of bacterial DNA input (16,125 and 1000 pg) and PCR cycles (25, 30 and 35 cycles) was tested.

None of the tested conditions resulted in significant differences in both alpha and beta diversity measures (Supplementary Fig. 4). However, analysis of the sequencing depth showed that, while the different conditions did not influence the amount of reads in the donor samples (mean = 64437, stdev = 30753), these had an impact on the negative controls (included during the DNA extraction step). A higher number of PCR cycles resulted in a higher amount of contaminant reads in the negative extraction controls (average number of reads detected in blank samples of 116, 4045 and 36044 for 25, 30 and 35 cycles respectively). For 35 cycles, the number of reads detected in negative controls were even comparable to those observed for true samples (Fig. 10).

**Figure 10.**
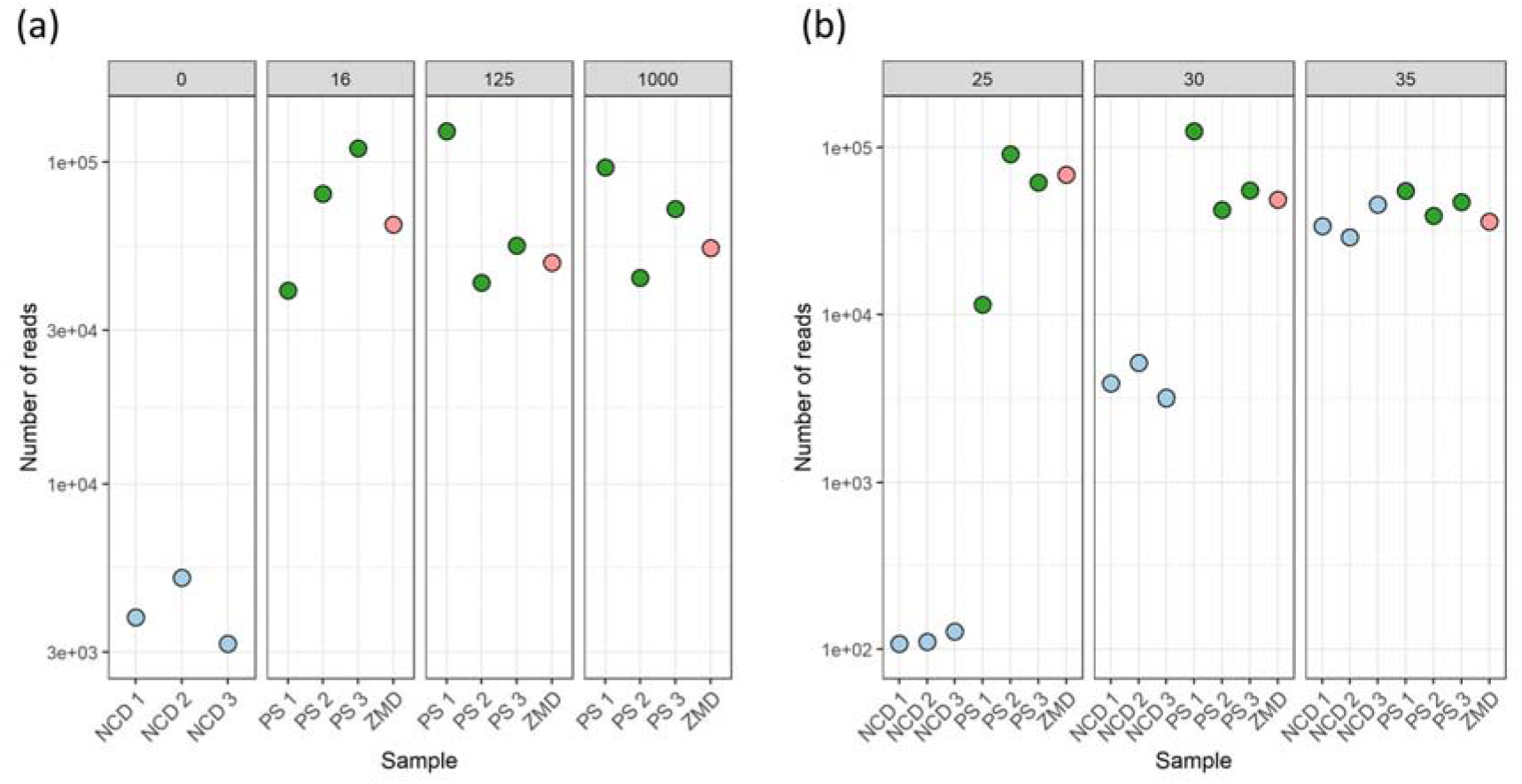
Sequenced reads of 3 participant samples (PS), 3 blanks (B) and 1 positive control (ZMC). The effect of different bacterial input (a) and PCR cycles (b) during the 16S rRNA gene V4 region PCR on the number of reads sequenced.

To identify the contaminant reads introduced in the different steps, we used the decontam package. Prevalence-based identification detected 24 contaminant ASV’s when analyzing all negative extraction controls (Table 2). Negative extraction controls (B) prepared with 35 cycles during amplification showed the highest number of reads for all (24/24 taxa detected, average number of reads 14450), where *Delftia* showed to be the main contaminating genus. When using 30 cycles we observed a reduction of read counts for all contaminants (23/24 taxa detected, average number of reads 1459). Samples prepared with 25 cycles showed the lowest number of contaminating sequences (9/24 taxa, average number of reads 45). Contaminants from the *Comamonadaceae, Ralstonia* and *Mesorhizobium* genera were also found in donor 1 (using 30 and 35 cycles) and a mock community sample (using 30 cycles). None of the contaminant sequences were identified in the true samples when amplified with 25 PCR cycles.

**Table 2.** Table of contaminant ASV’s as detected by the decontam package using the prevalence filtering.

| Contaminants in negative extraction controls (absolute reads) |  |  |  |
| --- | --- | --- | --- |
| ASV | 25 cycles | 30 cycles | 35 cycles |
| *Delftia* | 27 | 627 | 5621 |
| *Paenibacillus* | 6 | 84 | 3233 |
| *Comamonadaceae* | 8 | 245 | 1829 |
| *Massilia* | 3 | 77 | 256 |
| *Mesorhizobium* |  | 71 | 875 |
| *Ralstonia* |  | 83 | 392 |
| *Burkholderia-Caballeronia-Paraburkholderia* |  | 57 | 411 |
| *Ralstonia* |  | 46 | 228 |
| *Methylobacterium-Methylorubrum* |  | 0 | 205 |
| *Hydrogenophilus* |  | 19 | 176 |
| *Anoxybacillus* |  | 9 | 156 |
| *Ruminococcus* | 1 | 33 | 138 |
| *Neisseria* |  | 10 | 159 |
| *Meiothermus* |  | 4 | 155 |
| *Sphingobacteriales* |  | 6 | 119 |
| *Bradyrhizobium* |  | 24 | 113 |
| *Kapabacteriales* |  | 11 | 112 |
| *Sphingobacteriales* |  | 5 | 84 |
| *Lawsonella* |  | 11 | 66 |
| *Bacteroides* |  | 11 | 50 |
| *Meiothermus* |  | 5 | 28 |
| *Corynebacterium* |  | 7 | 23 |
| *Brucella* |  | 8 | 11 |
| *Micrococcus* |  | 6 | 12 |
| **Sum** | **45** | **1459** | **14450** |

| Contaminants in participant samples (percentages) |  |  |  |
| --- | --- | --- | --- |
| ASV | 25 cycles | 30 cycles | 35 cycles |
| *Comamonadaceae* |  | 0,00032 | 0,00034 |
| *Ralstonia* |  | 0,00013 | 0,00014 |
| *Ralstonia* |  | 0,00003 |  |
| *Mesorhizobium* |  | 0,00007 |  |
| **Sum** | **0** | **0,00055** | **0,00048** |

| Contaminants in positive controls (percentages) |
| --- |

| ASV | 25 cycles | 30 cycles | 35 cycles |
| --- | --- | --- | --- |
| Comamonadaceae |  | 0,00008 |  |
| Sum | 0 | 0,00008 | 0 |

Furthermore, we observed that the mock community sample showed the highest correlation to the theoretical composition when using 125 pg input DNA and 25 PCR cycles (rho= 0,833). In mock communities, changes in the number of PCR cycles have a greater impact on the microbial composition than the DNA input, mostly driven by a higher proportion of *Escherichia coli* and *Salmonella enterica* (Fig. 11).

**Figure 11.**
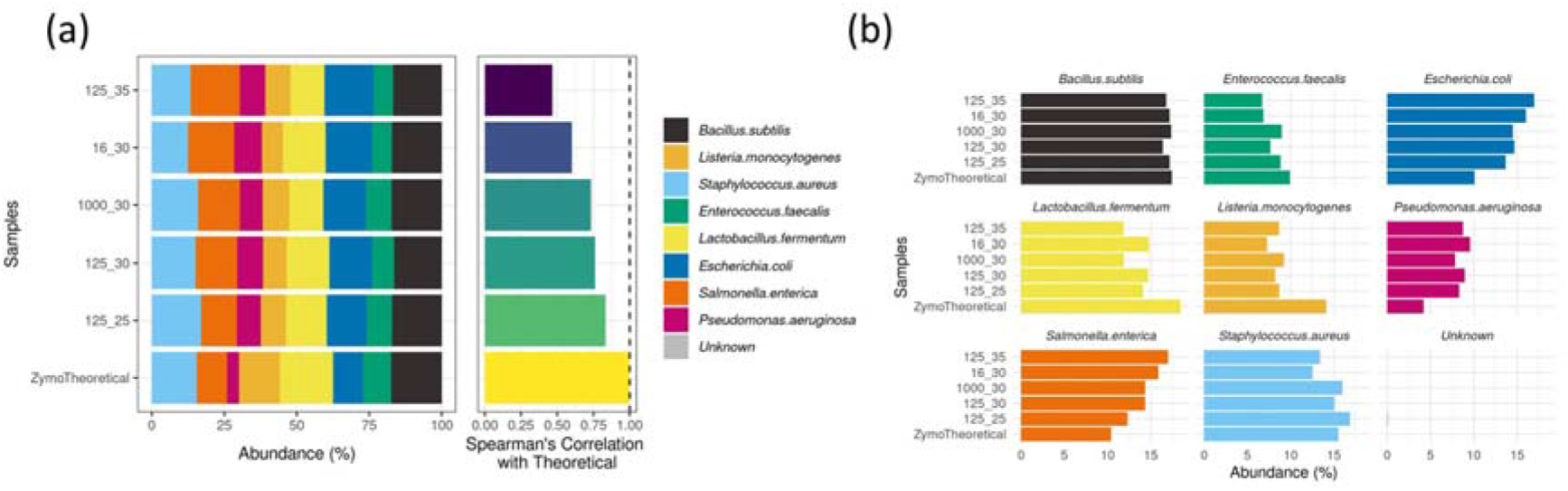
(a) Effect of PCR conditions on mock communities during library preparation compared to the theoretical composition using a Spearman’s correlation. (b) Barplots of the 8 bacterial strains included in the Zymo Mock sample

### Inter-run variation during sequencing

In our study, we included 30 mixed samples and 68 mock community samples to investigate the effect of sequencing in different runs, an approach generally used in large microbiome studies (Fig. 12). These samples were classified in 4 different groups: faecal samples extracted in different DNA extraction rounds (DE), DNA samples amplified with unique barcodes (UB) and DNA samples amplified with repeated barcodes (RB1 and RB2).

**Figure 12.**
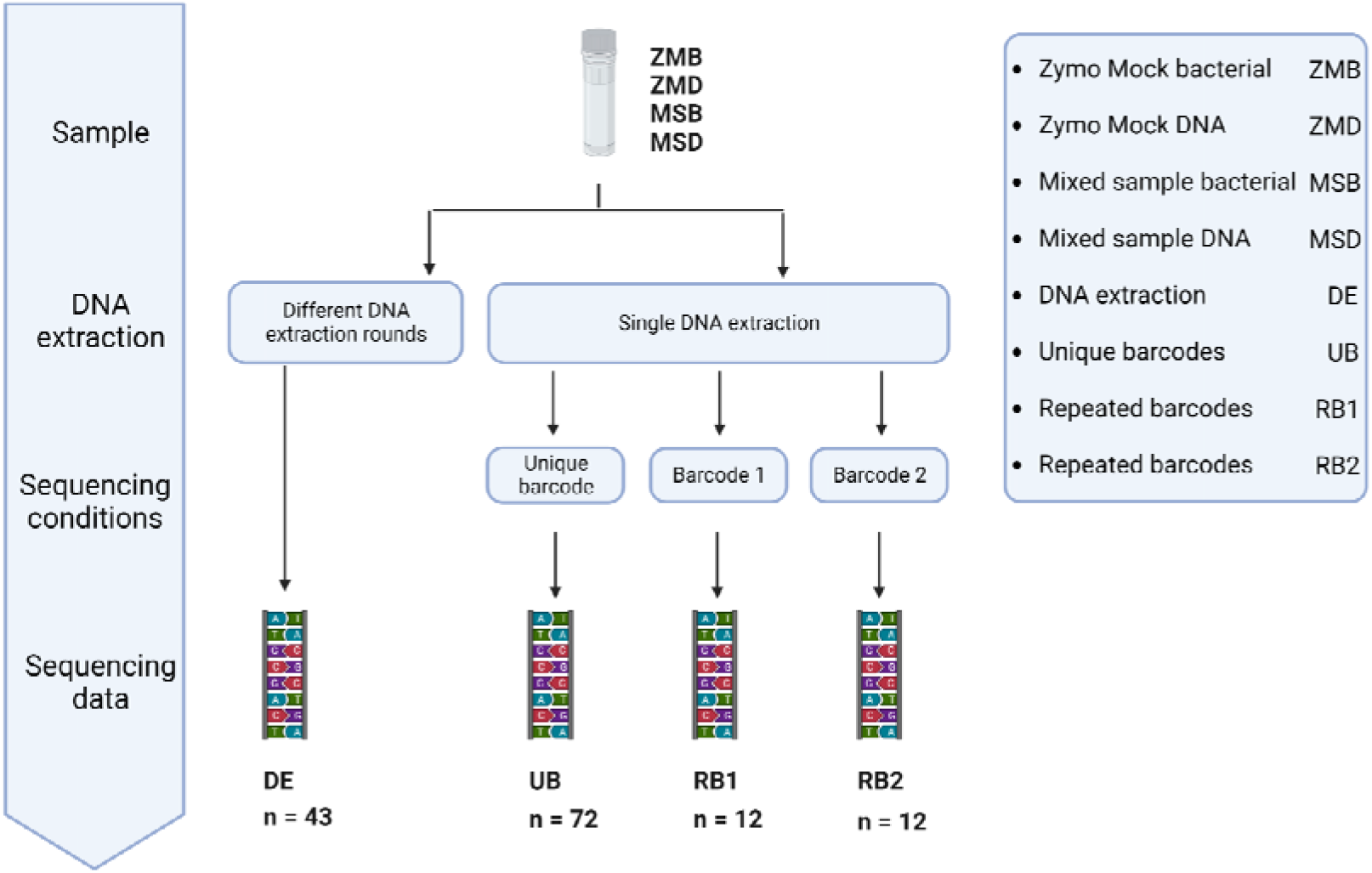
Overview of the conditions tested to measure the variation in large microbiome studies introduced during different DNA extraction rounds, multiple sequencing runs and the use of barcodes as a unique identifier.

*We* calculated the group divergence, a measure to quantify microbiota heterogeneity within a given sample set with respect to a reference, using the Bray-Curtis dissimilarity method. Although none of the conditions tested were significantly different, mock samples sequenced with the same barcodes (RB1 and RB2) showed the lowest divergence (Fig. 13a). Significant differences were observed when looking at the more complex mixed samples, showing the largest heterogeneity in the samples extracted in different DNA extraction rounds (DE) compared to the samples extracted in one run (UB). A significantly higher divergence was also found in samples sequenced with different barcodes (UB) when compared to those using the same barcode (RB1 or RB2) (Fig. 13b).

**Figure 13.**
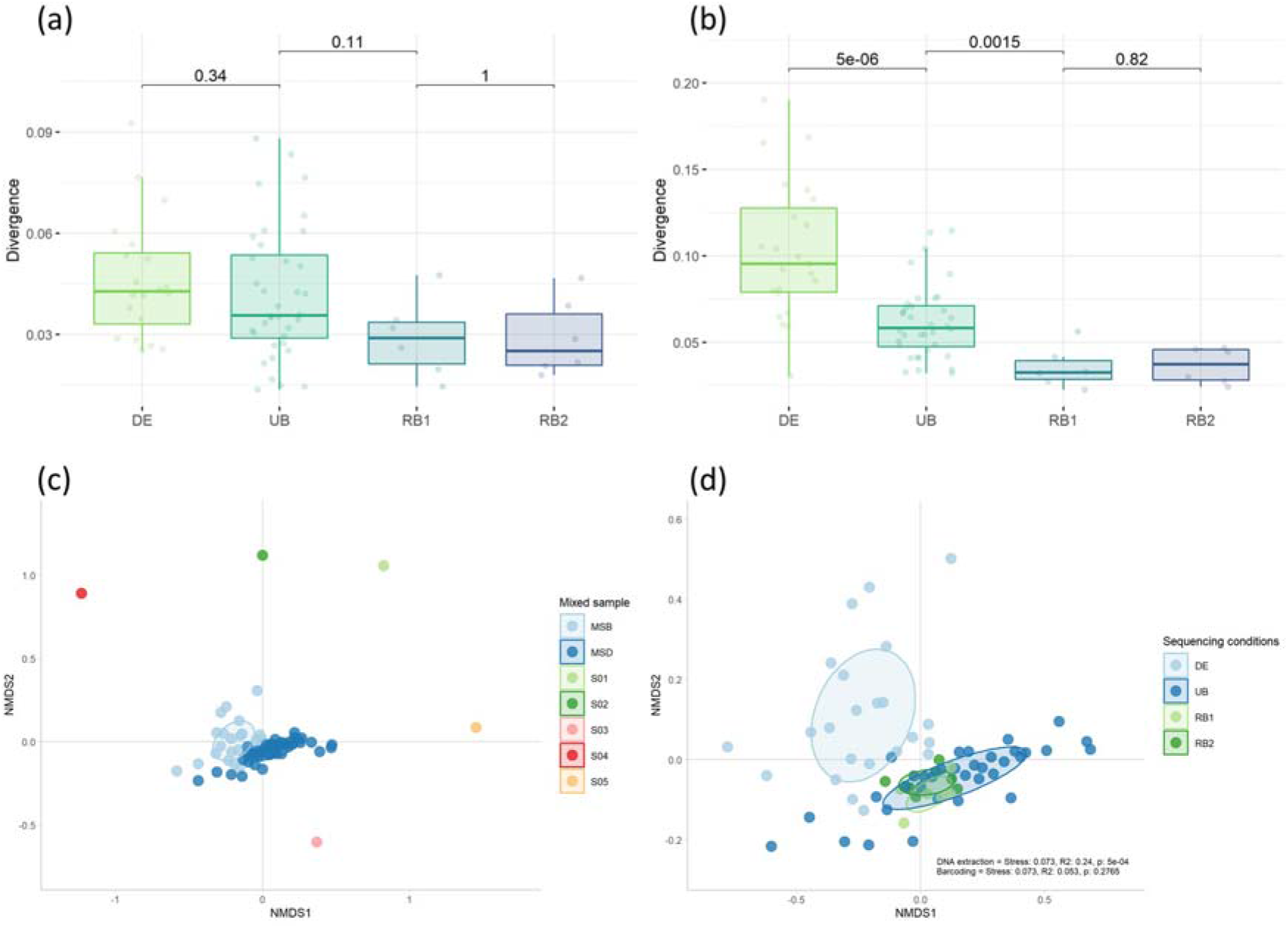
Divergence of (a) Mock samples and (b) mixed samples when comparing the effect of DNA extraction rounds and the use of different barcodes on the heterogeneity within the set of samples. (c) Non-metric multidimensional scaling (NMDS) plot of the 5 donor samples (S01 to S05) used to generate the mixed sample used as positive control in DNA extraction (DE) and sequencing runs (UB, RB1 and RB2) (n=76). (d) NMDS plot of the 30 mixed samples sequenced: faecal samples extracted in different DNA extraction rounds (DE), DNA samples amplified with unique barcodes (UB), DNA samples amplified with the same barcodes (RB1 and RB2).

*We* analysed the overall bacterial communities using a non-metric multidimensional scaling (nMDS) approach (Fig. 13c, d). We observed largest variation within the DE group, indicating that, from the conditions tested, DNA extraction is the most impactful step during sample preparation for sequencing. Using different barcodes (UB) during sequencing has a modest effect on the microbial composition when compared to the results obtained with the use of the same barcode (groups RB1 and RB2).

## Discussion

Biases introduced by methodological differences have been shown to influence microbiome profiles [13, 33, 34], making it difficult to compare results between studies and research groups. To overcome this challenge, numerous studies have been done to establish a standardized way of setting up microbiome studies [11, 35, 36], including a comprehensive checklist (STORMS) developed by a multidisciplinary team of researchers, which can be used as a guide towards a concise and complete reporting of microbiome studies [12]. Despite the importance of these validated protocols, in a field that is continuously evolving, it is essential to keep innovating methods to improve our current standards. Furthermore, with microbiome studies increasing in number and size, study designs and methodology need to be optimized on a larger scale to improve accessibility for underrepresented geographical locations. In this work, we aimed to contribute to the existing knowledge on the impact that different steps during a microbiome study can have on the overall microbiota diversity and composition, and to provide more insight into the biases introduced during the entire workflow and how to control for these.

Sample collection using the standard method (i.e. immediately freezing upon sampling and transported to the laboratory maintaining the cold-chain) is not always feasible [13, 37–39]. Collecting and transporting samples at room temperature however promotes the overgrowth of facultative anaerobes and potential degradation of nucleic acids by breakdown of strict anaerobes. To prevent this, several methods including different stabilization buffers have been developed in the recent years [38, 40, 41]. In line with the results reported by Roesch et al. [42], we found a higher amount of *Enterobacteriaceae* in samples stored at RT for 3 – 5 days, when compared to the same sample when directly frozen. This was largely prevented by the use of the Omnigene gut and the Zymo research DNA stabilization systems. Both stabilization methods, RT-ZB and RT-OB, resulted in microbiome profiles with lower proportions of *Actinobacteria*, a finding that has been reported before in studies comparing the Omnigene Gut to other collection methods [14, 37, 39]. Considering the fact that this decrease in *Actinobacteria* is not observed in the samples stored at RT without buffer, this is most likely to be an effect of storage in stabilization buffer. Furthermore, in samples stored at RT, including those stabilized using the Omnigene gut system, we observed an underrepresentation of *Firmicutes*, including the obligately anaerobic *Lachnospiraceae* (RT-ZC and RT-OB samples) and *Oscillospiraceae* (RT-ZC samples), indicating a suboptimal preservation of strict anaerobic bacteria. This difference was not observed with the Zymo Research system, indicating this to be a better preservation for *Firmicutes*. In previous studies, the Zymo research system was found to be suitable for preservation of chicken faeces [43] and soil samples [44]. From our results, we found the Zymo Research system to have the least differences when compared to the immediately frozen samples, although results of the abundance of *Actinobacteria* should be interpreted with caution, because differences could be induced by the method for sample storage.

Differences in DNA extraction methods have been shown to have the highest impact on the microbial composition, where mechanically disruption of the cells has proven to give the most reliable outcomes [10, 45, 46]. As expected from previous literature, we found a higher DNA yield in the bead-beated samples without stabilization buffer, suggesting this to be a more effective way of cell disruption compared to enzymatical lysis. However, this difference in DNA yield was not observed in the faecal samples extracted in the presence of stabilizing buffer (RT-ZB), indicating that Proteinase K treatment is more effective in faecal samples that are previously diluted and homogenised. However, although DNA extraction with enzymatical lysis showed similar yield compared to mechanical lysis, we did observe a significant difference in the compositional profiles. Mock community samples showed highest correlation of the bead-beated samples when compared to the theoretical composition. Mock samples lysed enzymatically showed an underrepresentation of *Enterococcus faecalis* and *Listeria monocytogenes*, both gram positive bacteria which are known to be more difficult to break [47]. In our study to improve reproducibility, we automated the lysate purification after cell disruption by using a Maxwell RSC system [48], comparing the Maxwell RSC Blood kit (traditionally used) and the newly developed Maxwell RSC Faecal kit. Both methods resulted in very comparable results with modest to no effect on diversity measures and compositional profiles. Although differences are very minor, when compared to the theoretical mock community composition, the Maxwell RSC Faecal kit showed a higher correlation, making this the preferable method for this sample type.

We observed that a higher number of PCR cycles lead to accumulation of chimera’s, point mutations and artifacts, as previously shown [49, 50]. Our data confirmed these observations showing an increase in contaminant ASV’s in the negative controls using more PCR cycles. However, most of these contaminants were no longer observed in the presence of faecal material or using a positive control (mixed sample or mock community), suggesting that this effect is mainly problematic during preparation of low biomass samples [18, 51]. Overall from our observations, we propose ~125 pg input DNA and 25 PCR cycles as optimal parameters during library preparation for human faecal samples when using the methods evaluated in this study. In cases were 25 cycles are not sufficient to obtain enough material (such as low biomass samples), extensive analyses and inclusion of negative controls should be exercised.

To track biases introduced in the various steps for library preparation and sequencing within a microbiome study, it is necessary to include repeated, complex, technical controls. Our data showed no significant differences in the heterogeneity measured in the different control groups using the commercially available community standard (ZMB or ZMD). Differences in factors such as GC content or presence of specific taxa in databases can influence the sequencing results, suggesting that mock communities alone are not sufficient to control for quality in faecal microbiome samples [52, 53]. We observed an effect when using the more complex mixed sample (MSB), although the limitation here is the unknown composition. However, this can serve as a technical replicate with properties similar to the true samples between multiple batches of sample processing. We found the highest heterogeneity in the DE group followed by the UB group, indicating that both DNA extraction and the use of different Illumina barcodes have an impact during library preparation. The use of 1 unique barcode resulted in a low heterogeneity in the samples, even though these were sequenced in different sequencing runs, indicating that there was no batch effect observed between the different runs.

Observations from our study may be subject to the low number of samples used for some of the analyses, i.e. to investigate the influence of the nucleic acid extraction and library preparation on the overall community structure. Furthermore, we are aware that we were not able to test all available methods and protocols, and we limited these to the commonly used approaches in our laboratory. Last, these results were obtained in the same laboratory, neglecting the effect of different equipment and laboratory environment. Collaborative studies between multiple laboratories is necessary to test the reproducibility and comparability of datasets from different studies. With novel tools continually emerging, future methodological work will always be needed to expand our current knowledge.

Standard guidelines to process microbiome studies will reduce technical variation across multiple studies, which will allow comparison of results between different research groups worldwide. Effective, accessible standardization will increase the available data to a broad range of diseases, ethnical backgrounds and geographic locations. Expanding microbiome data is necessary, as human microbiome research nowadays is dominated by research groups in highly developed countries, neglecting most of the world’s population [54–56]. A more global prospective will harness the full potential of the microbiome to use for targeted strategies of prevention, treatment, and maintenance of health.

## Supporting information

Supplementary results

