## Supplementary results for "Standardization of laboratory practices for the study of the human gut microbiome"

**Supplementary figures**

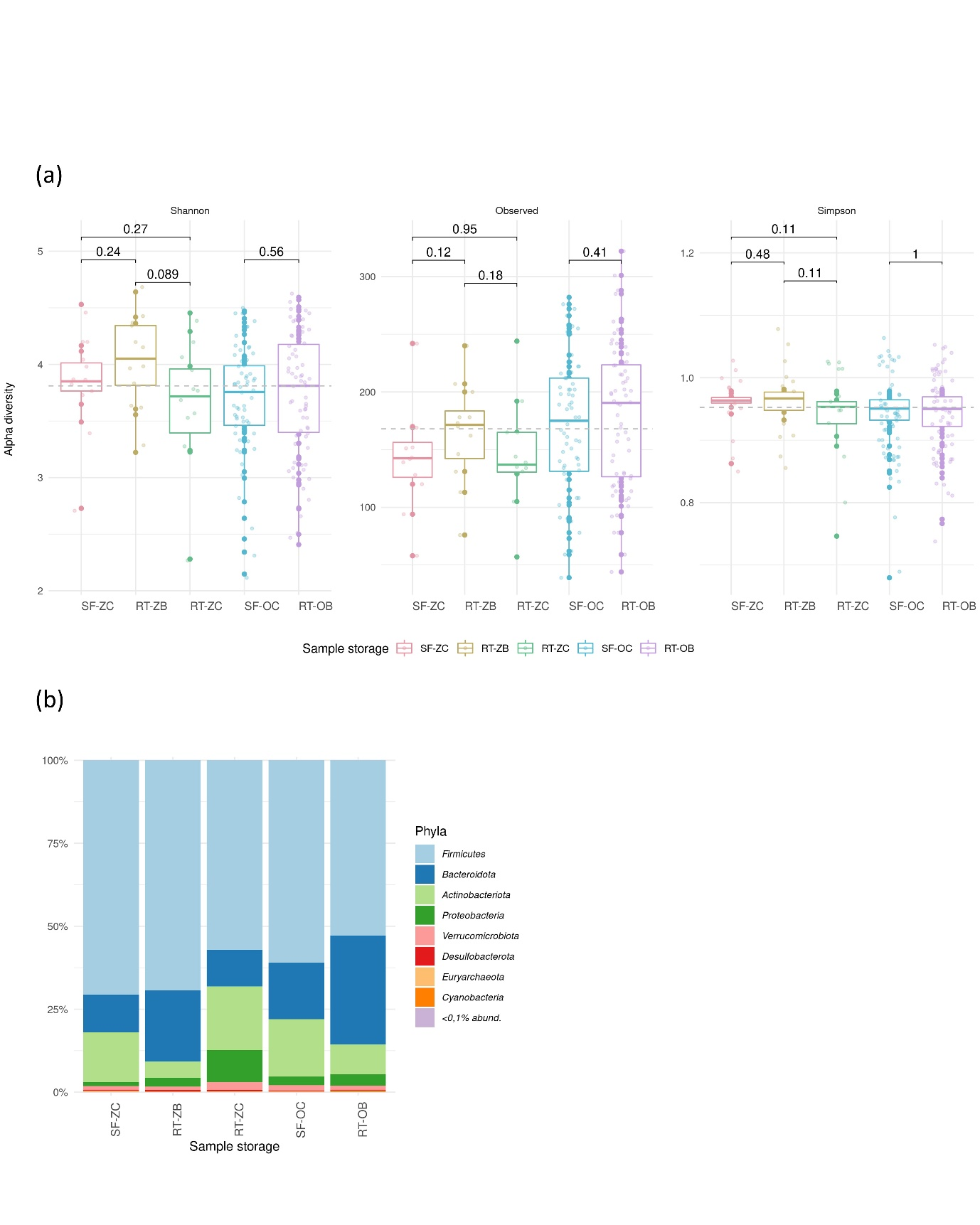

*Supplementary figure 1. Comparisons of sample collection and storage methods (a) Alpha diversity indices between different sample collection tubes and storage methods. (b) Bacterial composition at the phylum level, stacked bar chart shows the relative abundance of the phyla.*

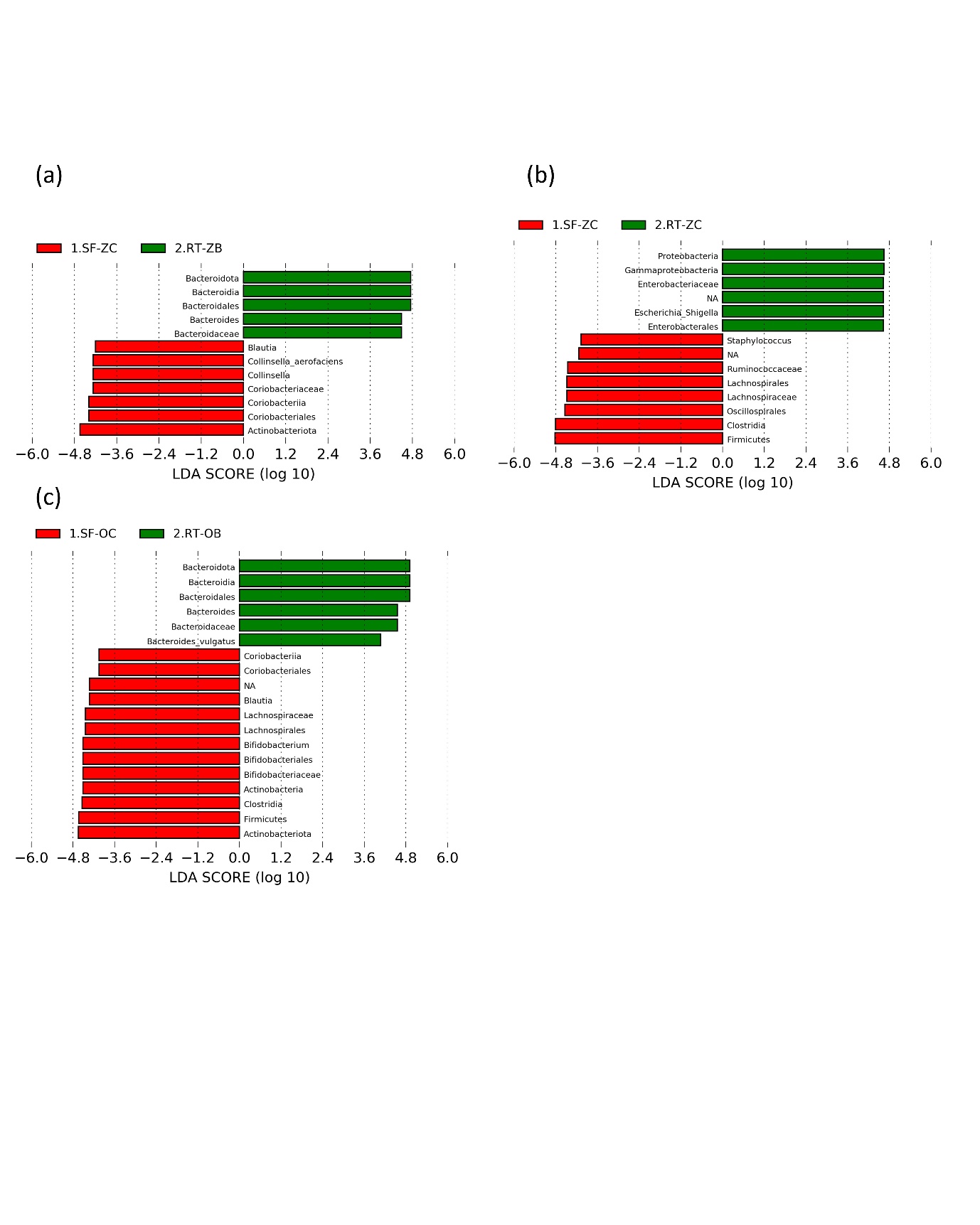
*Supplementary figure 2. LEfSe analysis showing the significantly distinguishing taxa between the different storage methods based on an LDA score > 4.0. Results are shown in bar graphs, showing the effect of storage at RT, with or without stabilization buffer (RT-ZB, RT-ZC, RT-OB) in green, compared to the immediately frozen control samples (SF-ZC, SF-OC) in red.*

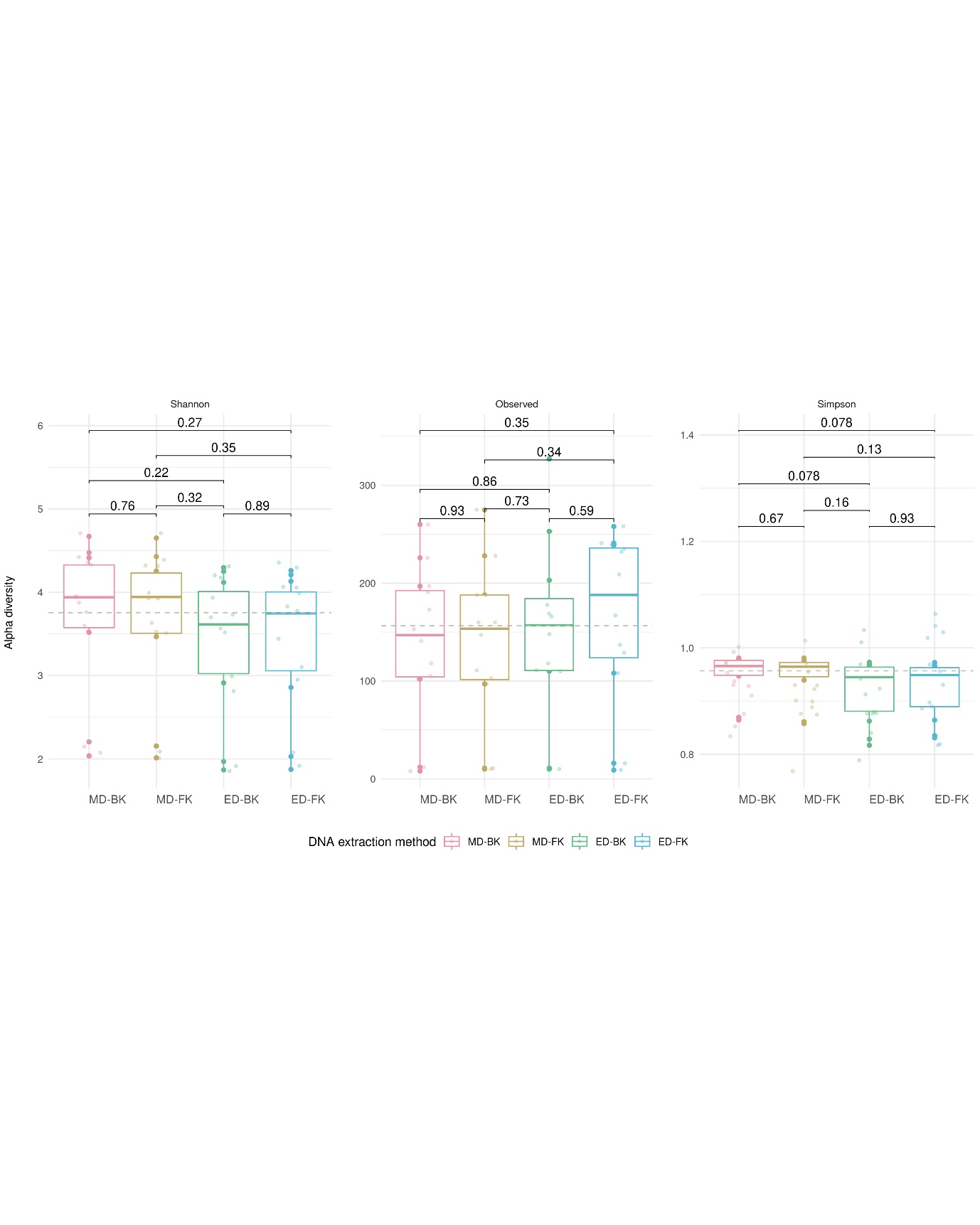

*Supplementary figure 3. Alpha and beta diversity measures. Comparison of Shannon index, Observed taxa and Simpson’s indices for the different extraction methods.*

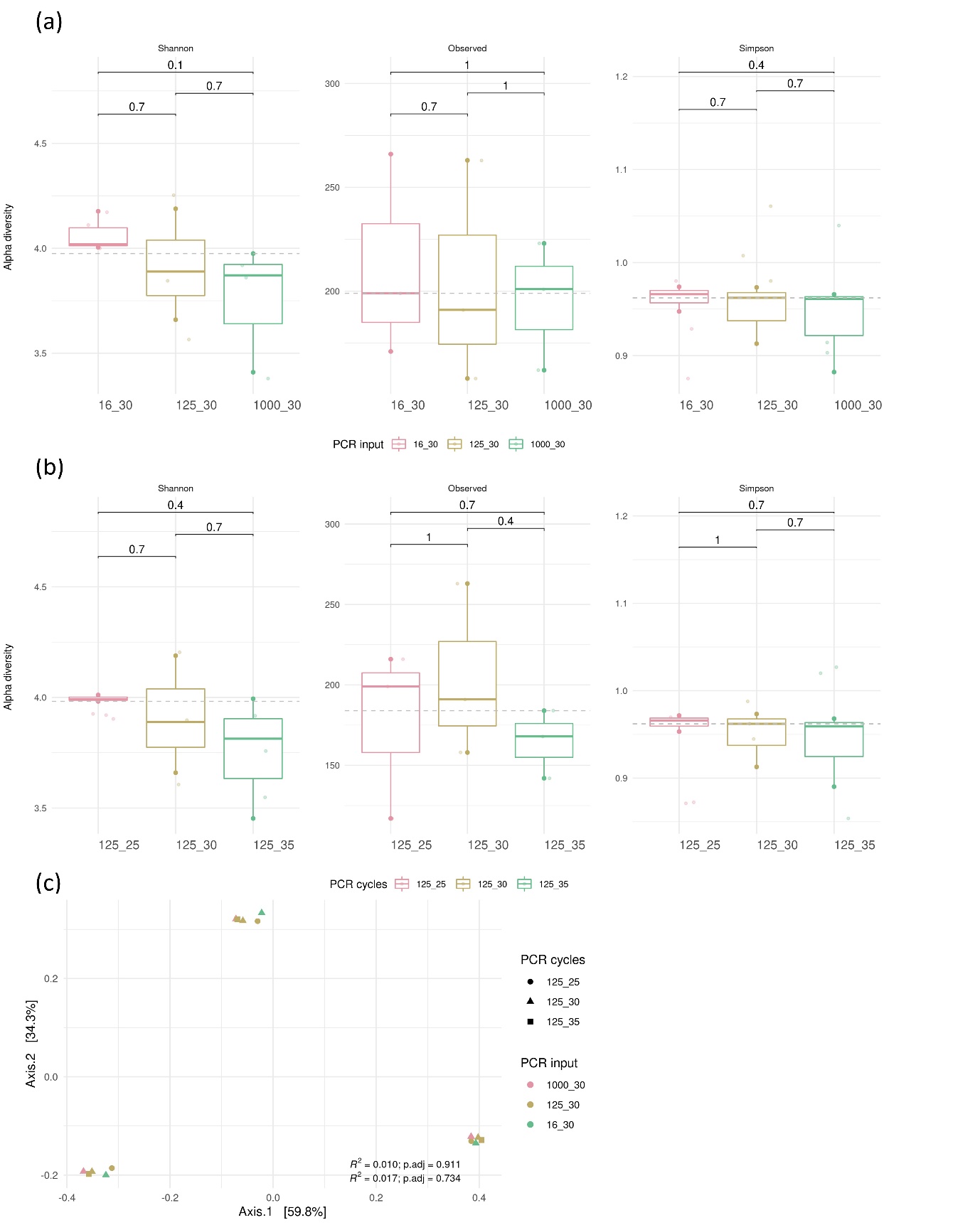

*Supplementary figure 4. Comparison of the different conditions tested by using alpha and beta diversity measures. (a) Shannon index, Observed taxa and Simpson’s indices were calculated for the different DNA input during amplification of the V4 region of the rRNA 16S gene.(b) Alpha diversity indexes for the different PCR cycles during V4 amplification. (c) Bray-Curtis distance in a PCoA ordination showing the difference in overall microbial composition of the different groups. perMANOVA results were displayed in the figure for the method for PCR cycles (R2= 0.010 ; p.adjusted= 0.911 ) and bacterial input (R2= 0.017; p.adjusted= 0.734)*
